# Single cell chromatin accessibility of the developmental cephalochordate

**DOI:** 10.1101/2020.03.17.994954

**Authors:** Dongsheng Chen, Zhen Huang, Xiangning Ding, Zaoxu Xu, Jixing Zhong, Langchao Liang, Luohao Xu, Chaochao Cai, Haoyu Wang, Jiaying Qiu, Jiacheng Zhu, Xiaoling Wang, Rong Xiang, Weiying Wu, Peiwen Ding, Feiyue Wang, Qikai Feng, Si Zhou, Yuting Yuan, Wendi Wu, Yanan Yan, Yitao Zhou, Duo Chen, Guang Li, Shida Zhu, Fang Chen, Qiujin Zhang, Jihong Wu, Xun Xu

**Affiliations:** BGI-shenzhen, Shenzhen 518083, China; Life Science College, Fujian Normal University; Key Laboratory of Special Marine Bio-resources Sustainable Utilization of Fujian Province, Fuzhou 350108, P. R. China; BGI Education Center, University of Chinese Academy of Sciences, Shenzhen 518083, China; Department of Neuroscience and Developmental Biology, University of Vienna; State Key Laboratory of Cellular Stress Biology, School of Life Sciences, Xiamen University, Xiangan District, Xiamen, Fujian 361102, China; MGI, BGI-Shenzhen, Shenzhen 518083, China; Eye Institute, Eye and ENT Hospital, Shanghai Medical College, Fudan University, Shanghai, China; Shanghai Key Laboratory of Visual Impairment and Restoration, Science and Technology Commission of Shanghai Municipality, Shanghai, China; Key Laboratory of Myopia (Fudan University), Chinese Academy of Medical Sciences, National Health Commission, China; State Key Laboratory of Medical Neurobiology, Institutes of Brain Science and Collaborative Innovation Center for Brain Science, Shanghai Medical College, Fudan University, Shanghai, China

## Abstract

The phylum chordata are composed of three groups: vertebrata, tunicate and cephalochordata. Single cell developmental atlas for typical species in vertebrata (mouse, zebrafish, western frog, worm) and tunicate (sea squirts) has been constructed recently. However, the single cell resolution atlas for lancelet, a living proxy of vertebrate ancestors, has not been achieved yet. Here, we profiled more than 57 thousand cells during the development of florida lancelet (*Branchiostoma floridae*), covering important processes including embryogenesis, organogenesis and metamorphosis. We identified stage and cluster specific regulatory elements. Additionally, we revealed the regulatory codes underlying functional specification and lineage commitment. Based on epigenetic features, we constructed the developmental trajectory for lancelet, elucidating how cell fates were established progressively. Overall, our study provides, by far, the first single cell regulatory landscape of amphioxus, which could help us to understand the heterogeneity and complexity of lancet development at single cell resolution and throw light upon the great transition from simple chordate ancestor to modern vertebrates with amazing diversity and endless forms.

## Introduction

Vertebrate evolved through two rounds of ancient whole genome duplications, followed by functional divergence in terms of regulatory circuits and gene expression patterns^1^. Lancelets are derived from an ancient chordate lineage, and belong to the cephalochordate clade^2^. As the sister group of vertebrates and tunicates, cephalochordate showed great similarity to fossil chordates from Cambrian in terms of morphology and thus have been considered as “living fossils”^3^, or living proxies for the chordate ancestor. As a slow-evolving chordate species, amphioxus is a good model to explore evolution history and provide insights about vertebrate origin and evolution. Amphioxus has been considered as a key phylogenetic model animal about vertebrate origin ^4^. In contrast to tunicates, cephalochordates maintained their basic body plans since Cambrian, therefore study on cephalochordates could probably provide valuable insights about the patterning mechanisms conserved among chordates. Yang et al. performed RNAseq for Branchiostome, covering 13 development stages, ranging from fertilized egg to adults. In total, they identified 3423 genes changed significantly during the development, including the increase of translation associated genes and decrease of cell cycle genes^5^. Marletaz et al. performed whole genome sequencing for Branchiostome lanceolatum and constructed genomic profiles including transcriptome, histone modification, DNA methylation and chromatin accessibility at bulk level. This study revealed high conversation of cis elements and gene expression patterns between vertebrates and amphioxus at earlier middle embryonic stage, supporting the hourglass model^6^.

Although previous studies revealed key components and regulatory mechanisms of lancelet development, they fail to detect rare cell populations and rapid changing pathway signals during animal development because of the diversity of cell types and complicated cellular interactions. To understand the genomic and epigenomic landscapes of animal development, single cell sequencing methods have been widely employed in developmental biology. The developmental process for mouse, zebrafish, clawed frog, worm and ascidian have been investigated at single cell resolution^7-11^, which revealed cell lineages and developmental trajectories. These studies elucidate the dynamic gene expression profiles during the development of rodent, fish, nematode and proto-vertebrate. However, a cellular and molecular taxonomy of more ancient species such as amphioxus is still missing. In this study, we profiled the chromatin accessibility and gene expression of *Branchiostoma floridae* at bulk levels covering 18 continuous developmental stages, including sperm, oocyte, zygote, 2 cell, 8 cell, 16 cell, 64 cell, blastula, early gastrulation (early-gas or EG), late gastrulation (late-gas, or LG), early neurula (early-neu, or EN), middle neurula (mid-neu, or MN), late neurula (late-neu, or LN), larva, early metamorphosis (early-meta, or EM), middle metamorphosis (mid-meta, or MM), late metamorphosis (late-meta, or LM). Meanwhile, we characterized the single cell chromatin accessibility from blastula to larva stage. Our study, for the first time, revealed the dynamic epigenetic of lancelet development at single cell revolution, which classified cells into different groups with distinct patterns of chromatin accessibility profiles. Moreover, we traced the turnover of cis regulatory elements of each cell lineage through the whole life cycle of *Branchiostoma floridae*.

## Results

### Profiling of cis regulatory elements during amphioxus development

To evaluate cis regulatory elements (CREs) dynamics of *Branchiostoma floridae* development, we performed assay for transposase-accessible chromatin with high throughput sequencing (ATACseq) for *Branchiostoma floridae* including 18 time points (Figure 1a). During the entire development period studied, we detected a total of 2,181,151 peaks after rigid filter criteria. To evaluate repeatability of each stage, we performed principal component analysis (PCA) (Figure S1a), which discriminated different development embryonic stages and indicated replicates of the same stage tended to share higher similarity. Genomic feature analysis (Figure S1b) showed more than 25% of peaks fell to the promoter regions. We found peaks located at promoter regions increased dramatically after early gastrulation stage and increased progressively from zygote to middle neurula stage, and decreased afterwards, which might indicate corresponding dynamics of regulatory elements during developmental stages. Interestingly neurula stage possesses the least number of peaks, but has the largest ratio of peaks within promoter regions.

**Fig 1:**
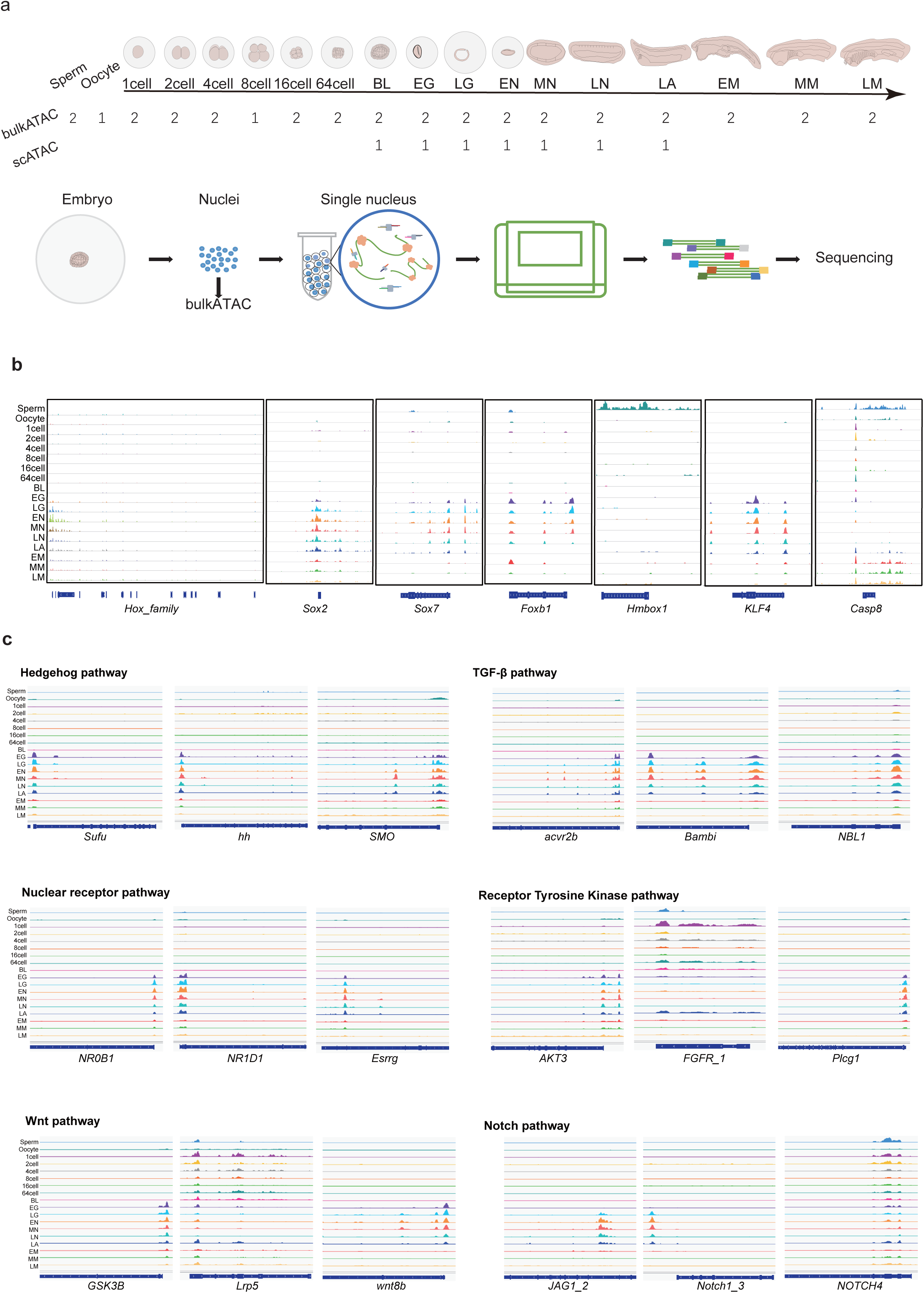
The accessible chromatin landscape and dynamics of B. floridae embryo. a) Schematic diagram showing brief experimental workflow and samples generated from different developmental stages, with number of biological replicates indicated for each sample type. Abbreviations and corresponding sample stages: BL. blastula, EG. early gastrulation, LG. late gastrulation, EN. early neurula, MN. middle neurula, LN. late neurula, LA. larva, EM. early metamorphosis, MM. middle metamorphosis and LM. late metamorphosis b) Genome browser view of a representative region showing the stage specific chromatin accessibility. c) Genome browser view of genes related to crucial developmental signal pathways

### CREs of master regulators and signaling pathway components were under dynamic regulation during amphioxus development

To pinpoint the profile of dynamic accessible regions during amphioxus development, we visualized ATACseq data of all time points using the integrative genomics viewer (IGV). *Hox* genes were reported to encode a family of transcriptional factors playing crucial roles in specifying body plans along the head-to-tail axis in various animals^12^. In *Branchiostoma floridae*, 15 *Hox* genes in total, existed in clusters on chromosome 17 within a 500kb range (Figure 1b). Based on our data, chromosome regions close to *Hox1_Bf, Hox2_Bf* and *Hox3_Bf* genes started to open dramatically from early gastrulation stage and maintained the epigenetic signals to the metamorphosis stages. Furthermore, signals of above three *Hox* gene regions were much stronger than that of *Hox4_Bf* to *Hox15_Bf*. Among *Hox* genes which harbored relatively less signal intensity, there were mainly four accessible patterns observed in different developmental stages. The first pattern was that signals were rather weak in sperm, oocyte and early embryonic stages, but sharply arose from early gastrulation stage and maintained till late metamorphosis stages. *Hox4_Bf* followed the first pattern. The second pattern was that regions opened in sperm, zygote and early gastrulation to late metamorphosis stages but were nearly closed in oocyte and early embryonic stages from 2-cell to blastula. *Hox5_Bf, Hox6_Bf, Hox7_Bf, Hox9_Bf, Hox10_Bf, Hox11_Bf, Hox13_Bf, Hox15_Bf* followed the second pattern. *Hox8_Bf* and *Hox12_Bf* followed the third pattern of which regions approximate to the genes seemed to be closed across all embryonic stages. The last pattern, which *Hox14_Bf* followed, was that peaks emerged in sperm, zygote, 2cell, early gastrulation, early neurula, early metamorphosis to late metamorphosis stages but were hardly seen in remaining stages.

Signals of peaks near *Sox2, Sox7* and *KLF4* genes emerged in early gastrulation stage, maintained till larva stage, and weakened afterward. *Sox2* was previously reported as evolutionally conserved and essential for embryo stem cells and neural progenitor cells pluripotency in various of species ^13^. *KLF4* could act both as activator and as repressor and was involved in regulating transcription, cell proliferation and differentiation, and maintaining stem cell population^14^. *Hmbox1* gene was near the open regions which were specific in sperm stage while *Casp8* gene tended to be near the open peaks in almost all stages except middle neurula and late neurula.

Cell to cell communication pathways are of great importance for the commitment of cell fates during amphioxus development ^15^. Here we investigated chromatin accessibilities of genes involved in crucial pathways during amphioxus development, namely Hedgehog, Notch, Nuclear Receptor (NR), Receptor Tyrosine Kinase (PTK), Transforming Growth Factor-β (TGF-β) and Wingless/Int (Wnt) pathways. (Figure 1c). Previous research reported amphioxus with mutation of *hh* gene showed deformities including a curled tail, no mouth etc., indicating a role of Hedgehog pathway in the left/right asymmetry control in amphioxus ^16^. Based on our data, CREs of *hh* gene were accessible from early gastrulation stage till larva stage, which was consistent with previous report that *hh* transcripts were first detected in the dorsal mesendoderm at the early gastrula stage^17^. *Lrp5* gene in Wnt pathway and *NOTCH4* gene in Notch pathway seemed to be ubiquitously accessible in almost all time points excepts in oocyte and late neurula stage. While *FGFR* gene in RTK pathway tended to be accessible during early embryo stages.

### Single-cell chromatin accessibility atlas of amphioxus development

We performed single cell ATACseq from samples ranging from gastrulation to larva stage, and obtained data for around 57 thousand cells passing quality control measures (Figure 1a). Specifically, the chromatin accessibility status of 11081, 7026, 6316, 9301, 14215, 9470 cells were detected from early gastrula, late gastrula, early neurula, middle neurula, late neurula and larva respectively. Low dimension visualization of the chromosome accessibility atlas at six stages revealed the increasing complexity during embryogenesis (Figure 2a). All 15 clusters identified by the similarity of epigenomic profile comprised no less than two adjacent sampled stages (Figure 2b), which confirm the comparability of a serials of samples. Next, we clustered cells at each time point and then classified chromosome accessibility of each cluster by a hierarchical clustering approach inferring the relevance of clusters. Then we annotated clusters by the accessibility of embryonic cell type markers of amphioxus.

**Fig 2:**
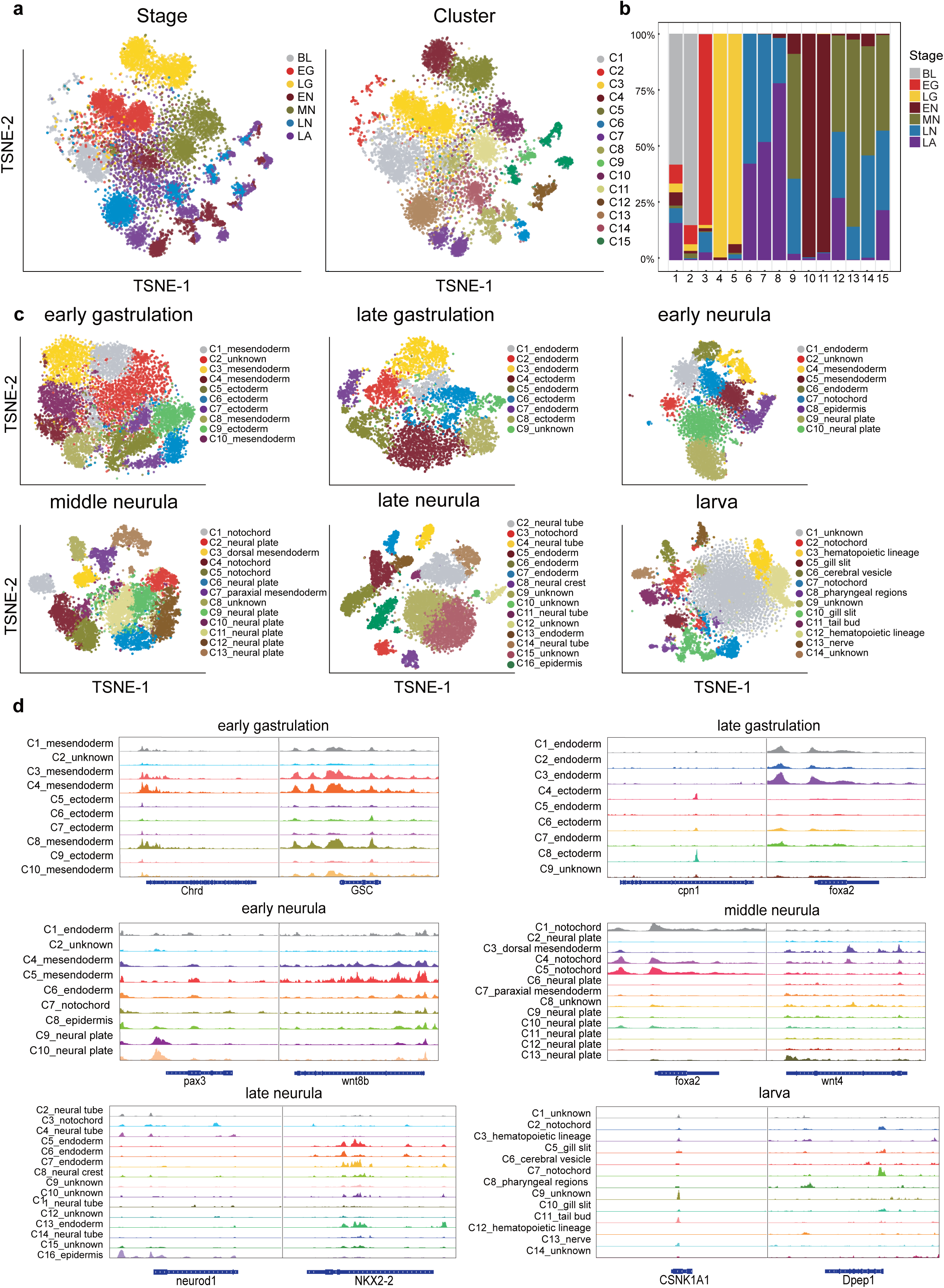
Single-cell chromatin accessibility atlas of B. floridae development. a) T-SNE map showing heterogeneity of cells from different developmental stages. Cells are colored by stages (left) and clusters (right) respectively. b) Proportion of cells from different stages in each cluster c) T-SNE projection of cells from indicated stages. Cells are colored by embryo lineage/cell types. d) Chromatin accessibility of lineage/cell type markers in indicated stages.

At early gastrulation stage of *branchiostoma floridae*, we identified 10 clusters corresponding to endoderm and ectoderm. EG1, EG3, EG4, EG8 and EG10 were defined as entoderm by the accessibility of *Foxa2*. EG4 and EG8 were specifically accessible at *DKK2/4*, the Wnt antagonist, which was the marker of the invaginating as the vegetal plate flattens^18^. EG5, EG6, EG7 and EG9 were identified as ectoderm by the accessibility of *Pax2*, and the neurol lineage marker *Mybl1*. At late gastrulation stage, we observed two distinct patterns corresponding to endoderm and ectoderm. LG1, LG2, LG3, LG5 and LG7 were identified as endodermal with high accessibility of canonical markers including *foxa2, sox7, gsc, afp etc*. LG4, LG6 and LG8 showed ectodermal characteristics due to the accessibility in *pax2* and *pax6*. In addition, neuronal lineage markers *nr6a1, pax3, sox2 etc*. were mainly accessible in LG4 and LG8. Taking into account of canonical Wnt/ß-catenin signaling which mediates cell fate decisions and cell movements during embryogenesis across metazoans, *CTNNB1* was accessible at EG5, EG6, EG7 and EG9 in early gastrulation stage. While in late gastrulation, *CTNNB1* was accessible concentratedly at LG8, inferring the cells located around the blastopore and it consisted with the finding that ß-catenin commonly localized at blastula stage, then was downregulated in mesendoderm at late gastrulation^15^.

At early neurula stage, we identified nine clusters corresponding to endoderm, mesendoderm, notochord, neural plate and epidermis. EN4 and EN5 was specifically accessible at *wnt8b*, which was the marker of mesendoderm. EN1 and EN6 were identified as endoderm by the accessibility of *Sox7*. EN9 and EN10 was identified neural plate as the specifically accessibility of *pax3* and *pax6*. EN8 was specifically accessible at *NEUROG3*, which was the marker of epidermis. EN7 was identified as notochord by the accessibility at *foxa2*. At middle neurula stage, MN2, MN6, MN10, MN11 and MN12 were accessible in *pax6, map2, gfap*, indicating ectodermal features. MN1, MN4 and MN5 was identified as endodermal lineage because of the accessibility in *foxa2, sox7, gsc* etc. Besides, we noticed notochord markers *shha, mnx1, dag1* showed accessibility in MN1, MN4 and MN5, suggesting the presumptive notochord characteristics. Additionally, MN3, MN7, MN9 and MN13 were identified as neuronal lineage as they were accessible in *neurod1, nefl, grin1* etc. At late neurula stage, LN5, LN6, LN7 and LN13 were accessible in *foxa2* and *nkx2-2*, indicating their endodermal features. LN2, LN11 and LN14 was identified as neural tube because of the accessibility in *sox2* and *hh*. LN4 was accessible at *phgdh* which is involved in neural tube development, was identified as neural tube as well. We noticed that notochord markers *shha* showed accessibility in LN3, suggesting the presumptive notochord characteristics. LN16 was accessible in *egfr* and *neurod1* which implied an epidermal feature, hence LN16 was annotated into neuroepithelial cell. Besides, LN8 was identified as neural crest cells due to the access of specific markers such as *neurog3* and *ascl1*.

In the larva stage, we identified 10 clusters corresponding to seven cell types: nerve cord, cerebral vesicle, notochord, pharyngeal region, gill slit, tail bud and hematopoietic lineage. LA8 were identified as pharyngeal regions by the accessibility in *NKX2-1*. LA2 and LA7 were accessible in *foxa2*, indicating the notochord feature, and LA13 was accessible in *Slc12a8*, which was the marker of nerve cord. LA6 was identified as cerebral vesicle with the accessibility in *pax6* and *neurog3*. We found that hematopoietic marker *vcan1, gblba* and *ncr1* showed specific accessibility in LA3 and LA12, suggesting the presumptive hematopoietic lineage characteristics. LA5 and LA10 were identified as gill slit with the accessibility in *pax9* and *tbx1*. LA11 showed the characteristics of tail bud due to the accessibility in *vasa*. Major organogenesis take place at larval stage. Interestingly, the pharyngeal region differentiates into distinct organs in larva with asymmetric patterning, the preoral pit and mouth on the left side and the endostyle and club-shaped gland on the right side. *Dkk2* was specifically accessible in LA8, which expressed at the margins of the mouth opening and revealed the left side position of LA8 in larva ^19^.

### Developmental trajectory of cephalochordate

We constructed putative trajectory based on the chromatin accessibility of canonical lineage markers. In global view, we observed distinct endoderm and ectoderm patterning in gastrula stage. Endoderm in late gastrulation, derived from early gastrulation, were later separated into mesendoderm, notochord and gut endoderm in early neurula. The gut endoderm lineage continued to differentiate into gill slit and foregut (pharyngeal region) in larva stage.

The development of notochord was also clearly captured. Following the first appearance of notochord cluster in early neurula (EN5), notochord clusters were identified in middle neurula (MN1), late neurula (LN15) and larva (LA2, LA7). *NTRK3* started to be accessible in late gastrula and remain accessible in following development. Similarly, the accessibility of another two nuclear receptors, *Nr2e3* and *hnf4a*, appeared in late gastrulation and increased gradually along the specification of notochord, finally peaking at larva stage.

Mesendoderm developed into hematopoietic lineage (LA5) in larva stage. We noticed that *FLT1* and *Flt4*, both of which belong to platelet-derived and vascular endothelial-growth factor receptors (PD/VEGFR receptors), showed distinct accessibility from late gastrulation stage. PD/VEGFR receptors have been linked with the formation of circulatory system in amphioxus, providing evidence for the validity of our trajectory.

In the separation point from late gastrulation endoderm to mesendoderm and notochord lineage in early neurula, we noticed that *NR5A2* was relatively more accessible in notochord (EN5) than in mesendoderm (EN1,7). The different accessibility pattern of *NR5A2* suggested the contribute of *NR5A2* to the segregation of mesendoderm and notochord.

As for ectoderm, we observed clear neural lineage in the trajectory. During the transition from late gastrulation to early neurula, ectoderm differentiate into two branches: epidermis and neural ectoderm. The epidermis layer persists along the embryo development. However, the neural ectoderm clusters then developed into neural plate (EN2,6,9-12), followed by the neural crest (LN3) and neural tube (LN4,9,11,14) in late neurula stage. subsequently in larva stage, neural tube clusters can be further developed into cerebral vesicle and nerve cord.

During the development course of neural lineage, we noticed the dynamic changes of several nuclear receptors. For example, *nr6a1* was highly accessible in gastrula stage (early gastrulation and late gastrulation) but became relatively closed in subsequent developmental stages. On the contrary, *NTRK3* and *Nr1h3* remained closed in gastrula stage but after which high accessibility were observed. We also found that *notch1* displayed a similar accessibility pattern to *NTRK3* and *Nr1h3*. Notch signaling has been reported to extensively involved the somites formation and segregation. Here we also suspected its role in neural lineage differentiation. It is worth noting that in larva stage, *NTRK3* was accessible in nerve cord but not in cerebral vesicle, suggesting a possible role of *NTRK3* to involved in the specification of nerve cord and cerebral vesicle.

### Transposable elements function as CREs associated with neural development during amphioxus development

Transposable elements (TEs) have been proposed to contribute to genetic complexity and provide genetic materials for the evolution of phenotype novelty. It has been reported that TEs are closely related to neural development. To investigate the functionality of TEs during lancelet development, we classified TEs into 67 families, and studied the biological process associated with each class. We found that PIF-Harbinger, LINE_I-Jockey, LINE_R2-Hero, DNA_Dada and TE_unknown family are associated with neural system development. Of particular, TE_DNA_PIF-Harbinger elements are closely related to “neurotransmitter metabolic process”, including the following genes: *PTX3, AGXT, PHGDH, SHMT1, PKD2, NOS1, TLR2*. TE_LINE_I-Jockey overlap with the regulatory elements of *GNPAT* related to ensheathment of neurons (Figure 4a). TE_LINE_R2-Hero element is located around *LRP4* gene related to regulation of neuromuscular junction development. Several elements in TE_unknown are related to neuron projection regeneration (*JAK2, LRP1, NDEL1, GRN, PTPRS, TNR, HGF, RTN4R, APOD, MATN2, TNC*) and regulation of neuron projection development (*PTK2, RET, LRP1, LRP4, CSMD3, NRP1, etc*.), positive regulation of neurogenesis (*RET, NOTCH1, LRP1, HES1, NRP1, NDEL1, etc*.), regulation of neuron projection regeneration (LRP1, NDEL1, GRN, PTPRS, TNR, HGF, RTN4R) and neural crest cell differentiation (*ALDH1A2, KBTBD8, RET, HES1, NRP1, LRP6, FN1, ALX1, KLHL12, NRP2*). In addition, an element in TE_DNA_Dada is related to neuroinflammatory response gene *FPR2* (Figure 4b).

**Fig 3:**
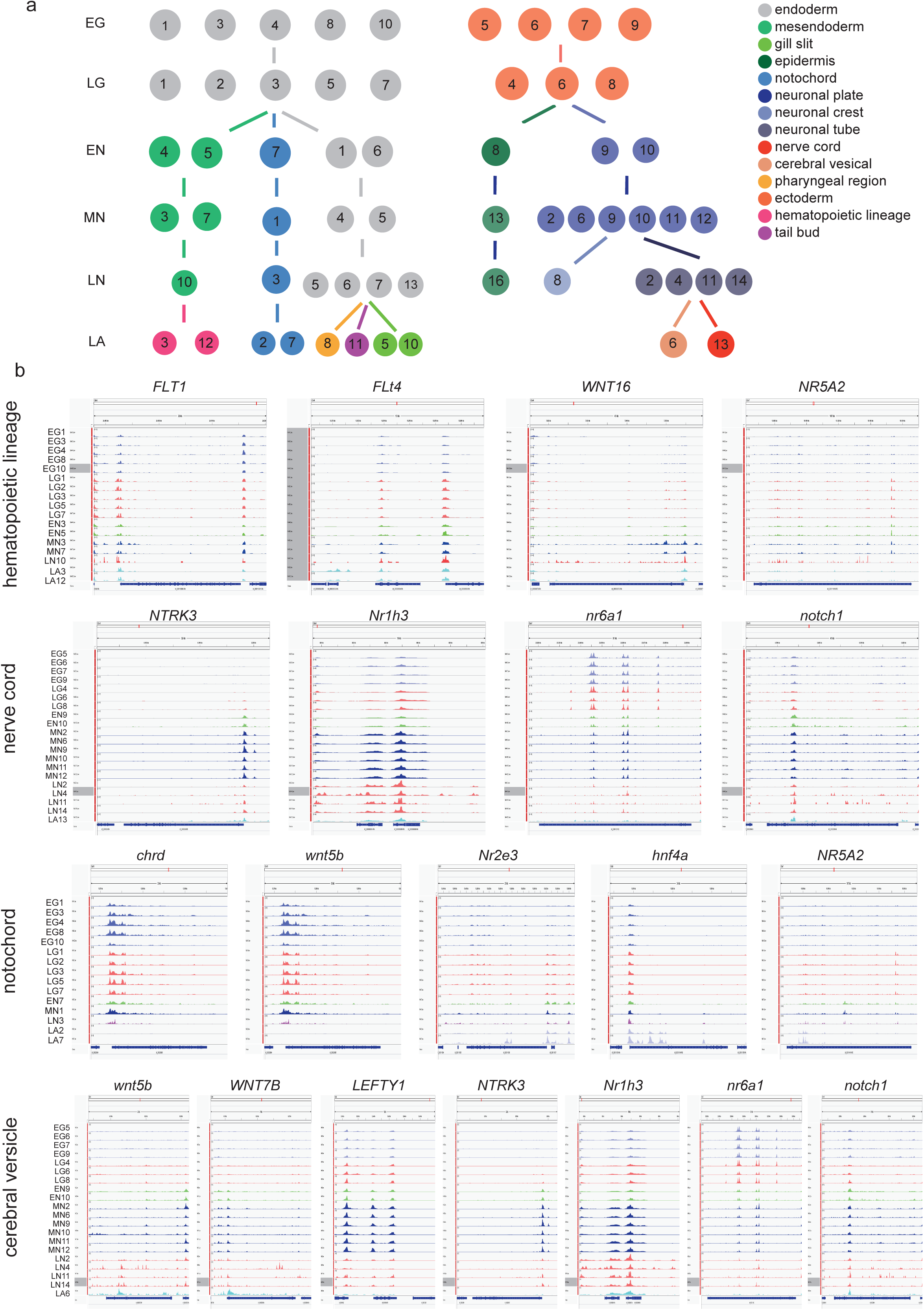
Epigenomic differentiation trajectories and genome browser views of genes. a) Differentiation trajectory of cells in each cluster along developmental stages, with stages indicated on the left. Numbers in circles represented clusters. b) Integrative genomics viewer of several function related genes with stages and clusters on the left side. EG1 represented cluster 1 of early gastrulation.

**Fig 4:**
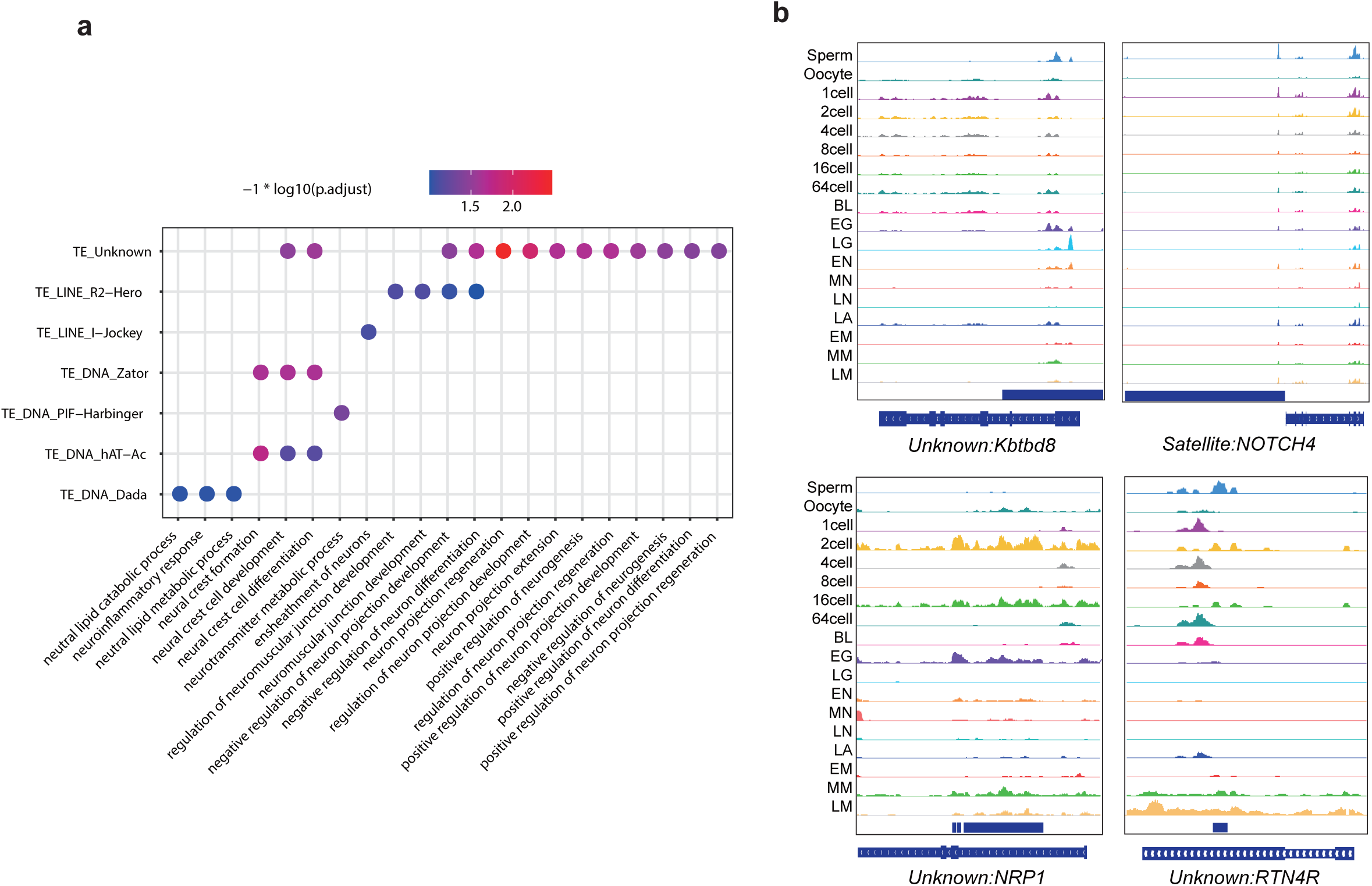
Transposable elements analysis. a) Bubble plot showing TE classes and enriched GO terms. b) Integrate genomic views of TEs and nearest genes.

## Discussion

Gene regulation is an important scientific question in developmental biology, as patterns are largely determined the spatial and temporal expression domains of developmental genes, which are decided by the dynamic interactions between TFs and CREs. CREs has been detected using experimental methods such as enhancer trapping, which is very time-consuming. We investigated the heterogeneity of cell populations of amphioxus and performed cross-stage and cross-cluster comparisons and revealed dynamic regulatory elements during several critical biological processes including gastrulation, neurulation and metamorphosis. Furthermore, we traced the chromatin states of key regulators controlling body plans and neural system development and inspected the dynamics of chromatin accessibility of components in WNT, TGF-β, RTK, nuclear receptor and Notch signaling pathways along developmental trajectory. Overall, we generated the first single cell epigenetic atlas for the developmental *Branchiostoma floridae*, a basal chordate lineage of particular value for studying the evolution and development of vertebrate. These results could be valuable for understanding the origin and evolution of chordate and throw light upon the genetic and epigenetic mechanism underling phenotype novelty of chordate during evolution.

## Methods

### Animal husbandry

Embryos were obtained by in vitro fertilization in filtered seawater and cultured at 25 °C.

### 10xATAC Sequencing Library preparation

Embryos were transferred to cold lysis buffer (Tris-HCl pH 7.4 10mM, NaCl 10mM, MgCl_2_ 3mM, Tween-20 0.1%, NP40 0.1%, Digitonin 0.01%, BSA 1%) and immediately pipetted 10X. Larva were homogenized in dounce homogenizer (SIGMA) 10X with lose pestle and 15X with tight pestle. Homogenization product was filtered through sterile 40-μm cell strainer (BD). Nuclei were centrifuged at 500 g for 5 min at 4 □ twice. First time nuclei were resuspended by chilled PBS (Gibco) with 0.04% BSA (BBI). Second time resuspend nuclei in diluted nuclei buffer (10X Genomics). Nuclei suspensions first were incubated in 10X Genomics transposition mix for 60 min, then transferred to Chromium Chip E to generate Gel bead-In-Emulsions according to Chromium Single Cell ATAC Reagent Kits User Guide Rev A (10X Genomics). After completed library construction step according to the Single Cell ATAC v1 workflow, library structure conversed by MGIEasy Universal Library Conversion Kit (App-A, MGI).

### Bulk ATAC-seq Sequencing Library preparation

For amphioxus embryos and larva, ATAC-seq was performed in two biological replicates following the original protocol^20,21^. Amphioxus embryos and larva were throwed in liquid nitrogen and frozen for 30 mins.

### Bulk ATAC-seq data processing

ATAC-seq was performed in more than two biological replicates at each developmental stage. ATAC-seq libraries were sequenced to produce an average of 78 million reads for amphioxus. Pair end raw data was generated by BGISEQ-500 sequencer and adapters were trimmed by Trimmomatic-0.39 with parameters “ILLUMINACLIP: adapter.fa:2:30:10 LEADING:3 TAINLING:3 SLIDINGWINDOW:4:15 MINLEN:36”. Bowtie2 was used for aligning with “--very-sensitive” and other parameters were left default. After aligning, duplicated reads were removed by MarkDuplicates function of Picard with “REMOVE_DUPLICATES=true”. Then reads with mapping quality below 30 or mapped to multi-positions of the genome were filtered. Reads remaining were used for downstream analysis. The model-based analysis mode of MACS2 was used for peak calling, and parameters were “-B -f BAMPE”. Only peaks with -logPvalue >=5 were kept for following steps. IDR (irreproducible discovery rate (IDR) was used for replicates merging. Gain and lost peaks were identified by BEDTOOLS intersect. R package of CHIPseeker was used for peak annotation. And Integrated Genomic Viewer was used for peak visualization. Read counts of every unique peaks were were calculated using summarizeOverlaps function of GenomicAlignments package with option “mode=‘Union’, inter.feature=TRUE, fragment=FALSE, singleEnd=TRUE”. This function also recepted a peak information which acted as a GenomicRanges object. We used rlogTransformation function of DESeq2 package to remove the batch effects between different repeats of same developmental stage. Dimensional reduction was applied using plotPCA function of BiocGenerics package and repeats were presented in 2-D plane. Then we removed the abnormal repeats according to the visual results. Gain and lost peak analysis between every two adjacent time points was performed by BEDTOOLS with “-v” parameter.

### TE analysis

Stage specific peaks were identified and merged by BEDTOOLS. TEs overlapped with merged peaks were classified based on their families and annotated to nearest genes by CHIPseeker. Genes of which TSSs were within the distance of 3k bp with TEs were used for gene ontology analysis. Position of TEs and peaks were visualized by IGV.

### Single cell ATAC-seq data analysis

Raw data sequenced by MGI2000 were split into reads and cell barcode using custom scripts. Reads were mapping to genome using snaptools with default parameters. To access the quality of each single cell, we calculated ATAC-seq quality metrics such as the fragment length distribution, transcription start site (TSS) enrichment and fraction of reads overlapping peaks were. We removed cell with total fragment less than 6000 and unique molecular identifiers (UMIs) less than 2500. Data was constructed into bins multiply by cells matrix. We remove the top 5% bins that overlap with invariant features. We sampled 40% to 20% of the total cells per stage from six developmental stages for following analysis. Then log-transformation, dimension reduction using diffusion map and clustering with K Nearest Neighbor (KNN) graph were applied using snapATAC. To identify the heterogeneity of clusters at each stage, we calculated the accessible signal of clusters using “bedGraphToBigWig”. Cell type annotation were based on cluster specific accessibility of marker genes visualized by IGV. We called peak by MACS2 with “-f BEDPE” and peaks with -logPvalue >=5 were kept for following DAR analysis.

### Sequencing and data availability

All 10xATAC sequence data was produced by MGISEQ-2000 in the China National GeneBank. ATAC-seq data was produced by BGISEQ-500 in the China National GeneBank. The data that support the findings of this study have been deposited in the CNSA (https://db.cngb.org/cnsa/) of CNGBdb with accession number CNP0000891.

## Acknowledgements

We are thankful to the production team of China National Gene Bank, Shenzhen, China.

## Figure legends

Fig S1

a) Principal component plots of accessible regions of each replicate with stages indicated.

b) Genomic distribution of peaks detected at each stage.

