## Supplementary figures and images for "Single cell chromatin accessibility of the developmental cephalochordate" (from another version of this preprint: version 1, 2020-03-18)

### Supplemental Figure 1

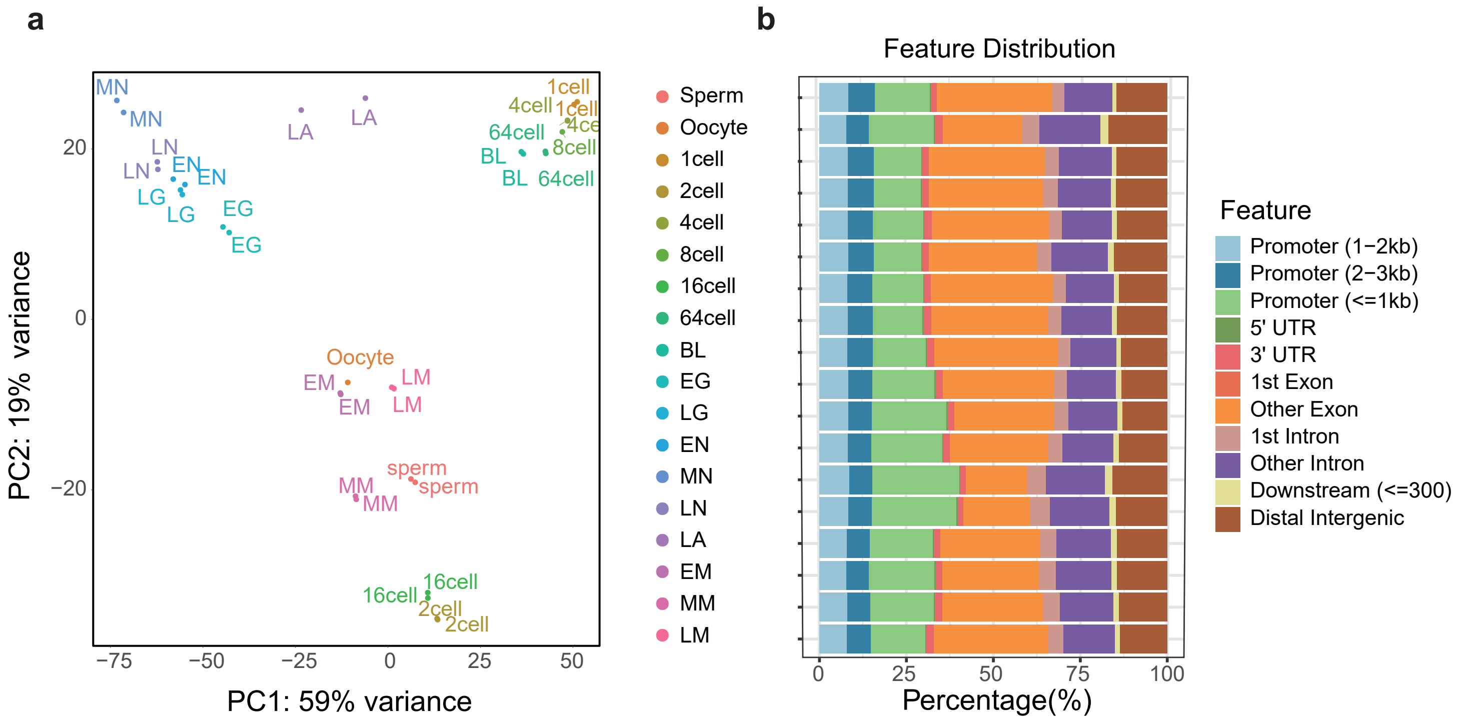
